# *CDeep3M* - Plug-and-Play cloud based deep learning for image segmentation of light, electron and X-ray microscopy

**DOI:** 10.1101/353425

**Authors:** Matthias G Haberl, Christopher Churas, Lucas Tindall, Daniela Boassa, Sebastien Phan, Eric A Bushong, Matthew Madany, Raffi Akay, Thomas J Deerinck, Steven T Peltier, Mark H Ellisman

## Abstract

As biological imaging datasets increase in size, deep neural networks are considered vital tools for efficient image segmentation. While a number of different network architectures have been developed for segmenting even the most challenging biological images, community access is still limited by the difficulty of setting up complex computational environments and processing pipelines, and the availability of compute resources. Here, we address these bottlenecks, providing a ready-to-use image segmentation solution for any lab, with a pre-configured, publicly available, cloud-based deep convolutional neural network on Amazon Web Services (AWS). We provide simple instructions for training and applying *CDeep3M* for segmentation of large and complex 2D and 3D microscopy datasets of diverse biomedical imaging modalities.

## Main

Biomedical imaging prospers as technical advances provide not only enhanced temporal^1^ and spatial resolution but also larger field of views^2^, and all at a steadily decreasing acquisition time. Altogether an abundant amount of potential information can be extracted from those very large and complex data volumes. In recent years, substantial progress has been made using machine learning algorithms for image segmentation^3^. Three-dimensional electron microscopy (EM) volumes, due to their extreme information content and increasing volume size^2,4^, are among the most challenging of segmentation problems and an area of intense interest for machine learning approaches. To this end, different architectures of deep neural networks^5–8^ show great promise towards one such challenge, the dense segmentation of neuronal processes spanning terabyte volumes of serial electron micrographs. However, generalized applicability of deep neural networks for biomedical image segmentation tasks is still limited and technical hurdles prevent the advances in speed and accuracy from reaching the mainstream of research applications. These limitations typically originate from the laborious steps required to recreate an environment that includes the numerous dependencies for each deep neural network. Further limitations arise from the scarcity of high-performance compute clusters and GPU nodes in individual laboratories, which are needed to process larger datasets within an acceptable timeframe. With the goals of improving reproducibility and to make deep learning algorithms available to the community, we built *CDeep3M* as a cloud based tool for image segmentation tasks, using the underlying architecture of a state-of-the-art deep learning convolutional neural network (CNN), DeepEM3D^6^, which was integrated in the Caffe deep learning framework^9^. While there is a growing number of deep-learning algorithms, we were attracted to the features offered by the CNN built in DeepEM3D^6^, as we recognized it to have advantages for our applications, where both cellular and subcellular features are of interest. Specifically, DeepEM3D is conceptually designed to be extremely broad in feature recognition with 18 million trainable parameters and three models trained in parallel on one, three and five consecutive image frames giving excellent results for - but not limited to - membrane segmentation^6^.

For *CDeep3M*, we modified all required components to make the CNN applicable for a wide range of segmentation tasks, permit processing of very large image volumes, and automate data processing. We also implemented a modular structure and created batch processing pipelines, for ease of use and to minimize idle time on the cloud instance. Lastly, we implemented steps to facilitate launching the most recent release of *CDeep3M* on Amazon Web Services (AWS). To reflect the now broad applicability of our implementation to data of multiple microscopy modalities (e.g., X-ray microscopy (XRM), light microscopy (LM) and EM), we named this toolkit *Deep3M.* To give users easy access and to eliminate configuration issues and hardware requisites, we further release a cloud-based version as *CDeep3M*. The publicly available AMI (Amazon Machine Image) of *CDeep3M* can be readily used for training the deep neural network on 2D or 3D image segmentation tasks. Using an AWS account gives any user the immediate ability to spin up a machine with *CDeep3M*, upload their training images and labels to generate their own trained model, and subsequently segment their datasets. To make this useful for researchers with varying levels of expertise, we minimized the number of sequential steps in *Deep3M* (**Fig. 1**), while still allowing for flexible use of the code.

Here we provide complete instructions for how to go from training images to performing segmentation using the deep neural network. Once the predicted segmentations are accomplished they can be post-processed either using an already available script (see supplemental material) or using standard image analysis and rendering tools (such as ImageJ, IMOD, Amira etc.) to group and count objects, mesh surfaces, or perform further analysis. We used *CDeep3M* for numerous image segmentation tasks in 2D and 3D, such as dense neurite segmentation, cellular organelle segmentation (nuclei, mitochondria, vesicles) or cell counting. We demonstrate the utility of this distribution of *CDeep3M* for LM, XRM, multi-tilt electron tomography (ET) and serial block face scanning electron microscopy (SBEM). Altogether this should facilitate the analysis of large scale imaging data and render *CDeep3M* a widely applicable tool for the biomedical community.

### Quick guide for using CDeep3M

While pre- and post-processing steps are less computationally intense, we include the necessary scripts in the pipeline on the same cloud environment for two reasons: first, to provide all the steps required to run the entire processing pipeline without knowledge in programming or the necessity to install or buy software and second, to minimize the traffic to and from the cloud environment. Data augmentation during pre-processing (Steps 1 and 3) increases the training data volume substantially (>16-100 fold). Also, the segmented results will be de-augmented in the post-processing (reducing it by a factor of up to a 100-fold). Typically, transfer of data will be more time consuming and therefore expensive than the added compute time on the cloud (<1min per 1000×1000×100 voxel). Steps 2 and 4 are run sequentially for 1frame (fm), 3fm and 5fm for 3D datasets for improved accuracy, whereas for 2D image segmentation only 1fm is run. Intense use of GPU occurs during steps 2 and 4 (training and prediction) requiring ^~^10GB of GPU memory at the current configuration.

### Data upload

It is necessary to upload training images and labels for steps 1-2 and raw data images for steps 3-4. All images can be uploaded as folders containing sequential tif or png images or a tif stack. Each will automatically be converted to the h5 file format during data augmentation. Using the png file format is recommended to minimize the data transfer. Training data consist of images and binary labels that use 0 as background and either 1 or 255 as positive label (see supplementary material for training data generation). A more detailed description of individual commands, together with additionally implemented scripts, is provided in the supplementary material. These scripts allow for more flexible use of the processing workflow, for example, to re-use previously trained models.

Briefly, the cloud resource is accessed using the secure shell command (ssh). The secure copy (scp) command is used to copy training images, labels, and image volumes to the cloud (and to copy predictions back). *CDeep3M* learning and processing consists of 5 steps (see **Fig. 1**):

Step 1: **Preprocessing / Data augmentation of training images and labels** PreprocessTraining ~/train/images/ ~/train/labels/ ~/augm_train
Step 2: **Training *Deep3M* CNN (steps 2 and 4 run automatically for 1fm, 3fm and 5fm)** runtraining.sh ~/augm_tr/ ~/train_out
Step 3: **Preprocessing / Data augmentation of images to segment** PreprocessImageData ~/images/ ~/aug_images/
Step 4: **Predict image segmentation (then performs data de-augmentation)** runprediction.sh ~/train_out/ ~/aug_images/ ~/predict_out
Step 5: **Ensemble prediction** EnsemblePredictions ~/predict_out/1fm ~/ predict_out/3fm ~/predict_out/5fm ./ensemble

**Figure 1:**
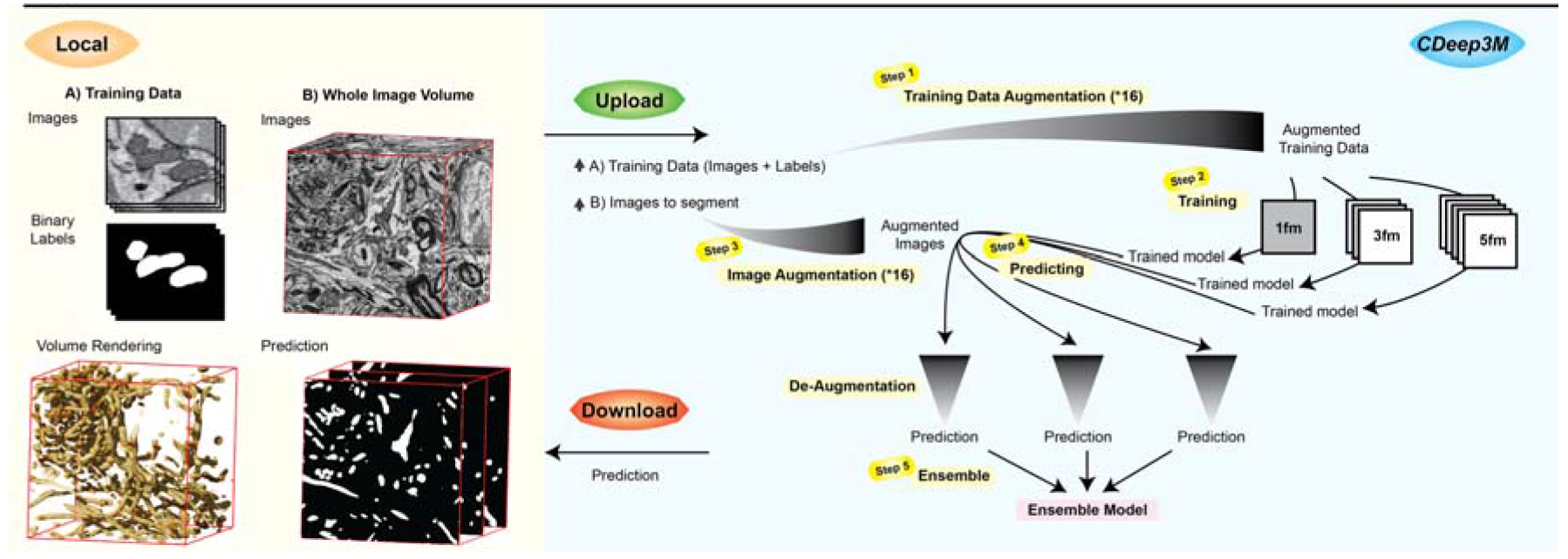
Image segmentation workflow with *CDeep3M*. Steps 1-2 are required to generate a new trained model based on training images and labels. For 3D models *CDeep3M* trains three different models (seeing 1 frame, seeing 3 frames and seeing 5 frames) that provide three predictions (Step 4), which are merged into a single ensemble model at the post-processing step (Step 5).

### Examples

When training *CDeep3M* for the segmentation of nuclei in XRM data of a hippocampal brain section, we established a cell density profile across the x-y-z directions of brain tissue prepared for EM (**Fig. 2a**). Training on fluorescence microscopy images of DAPI stained brain sections enabled us to distinguish neurons based on their chromatin pattern and distinguish one individual cell-type from other cells in the tissue (**Fig. 2b**). Multi-tilt electron tomography (ET) is a form of transmission EM used to achieve high-resolution 3D volumes of biological specimens. Here, we applied *CDeep3M* to high-pressure frozen brain tissue and were able to automatically annotate vesicles and membranes (**Fig. 2c**), which will aid the 3D segmentation of synapses and neuronal processes. We further used *CDeep3M* for the segmentation of intracellular constituents (nuclei, membranes and mitochondria) of the cell in a serial-block face scanning electron microscopy (SBEM) dataset (**Fig. 2d**). For our understanding of the role of intracellular organelles and alterations in diseases, parameters - such as the precise volume, the distribution, and fine details like contact points between organelles - will be of utmost importance. Therefore, we evaluated the performance (precision, recall and F1 value) of the underlying CNN based on predictions per pixel, which results in lower F1 values (compared to object detection) but is more representative to determine accurate segmentations and distribution of intracellular organelles. We found the segmentation accuracy of *CDeep3M* equals the one of human expert annotators (**Supplementary Fig. 1;** *CDeep3M:* F1 value 0.9039, precision 0.9052, recall 0.9027; humans: 0. 8827, precision 0.8993, recall: 0.8766).

**Figure 2:**
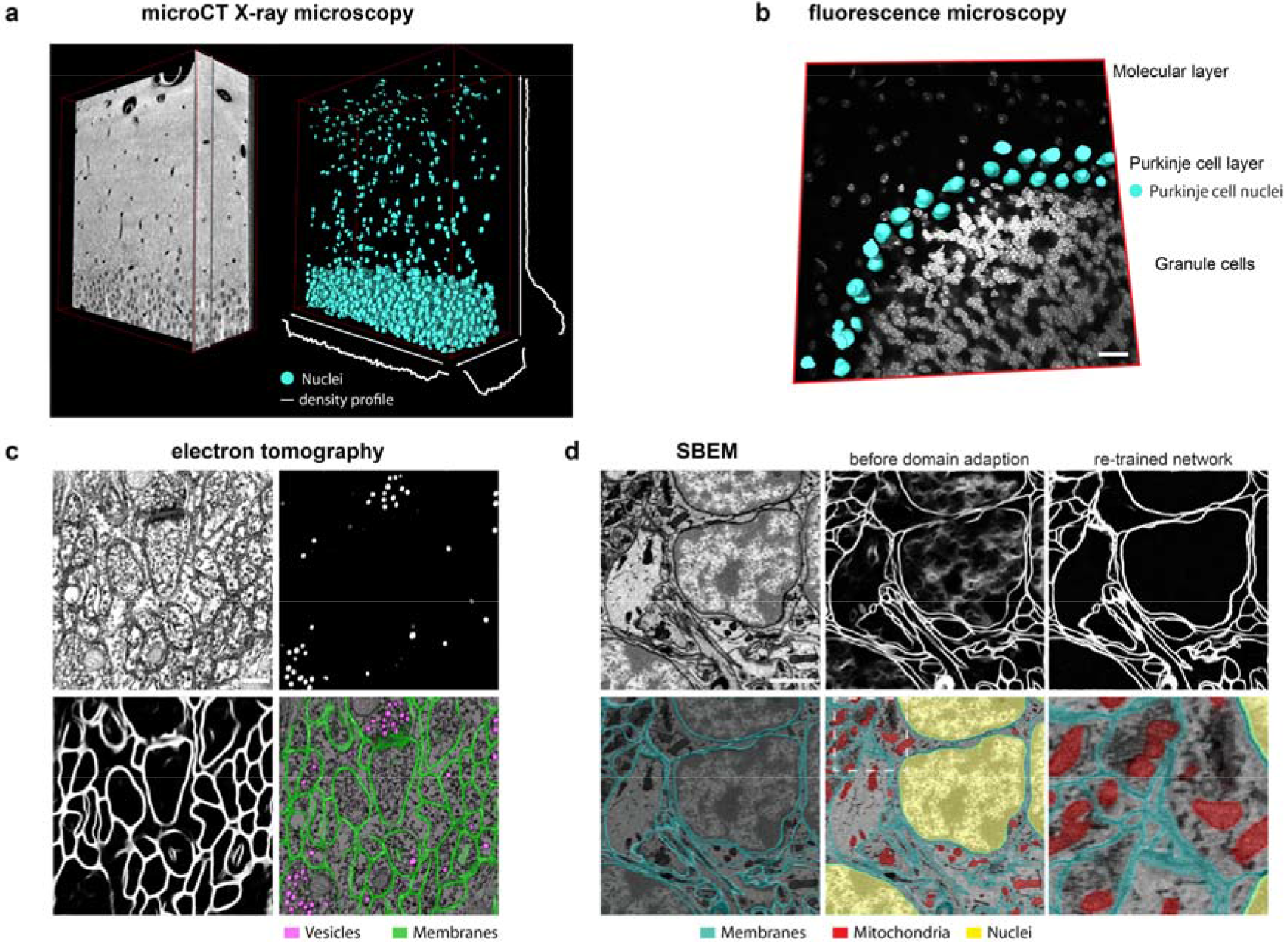
Multimodal image segmentation using *CDeep3M*. (**a**) Segmentation of nuclei in XRM volume of a 50μm brain slice containing the hippocampal CA1 area used for cell counting and establishing a cell density profile across x-y-z. (**b**) Segmentation of cell type specific DNA profile allows identification of Purkinje cells. Overlay of 3D surface mesh of nuclei on light microscopic image of DAPI-stained cerebellar brain section. Scale bar: 20μm. (**c**) Segmentation of vesicles and membranes on multi-tilt electron tomography of high-pressure frozen brain section. Scale bar: 200nm. (**d**) Upper row: SBEM micrograph (left) Scale bar: 1μm, segmentation using pre-trained model before (middle) and after domain adaption (right). Lower row: segmentation of membranes, mitochondria and nuclei overlaid on SBEM data

Since training is time consuming and costly, both to generate manual ground truth labeling and to train a completely naïve CNN, the re-use and refinement of previously trained neural networks (transfer learning) is of eminent interest. Generating manual training data for membrane annotation in SBEM is particularly laborious. To test our ability to re-use a pretrained model for a new dataset, we performed a type of transfer learning, domain adaption^10^. We first trained a model on the recognition of membranes with a published training dataset of a serial section SEM volume^11^ (similar to^6^) with a voxel size of 6×6×30 nm. We then applied the pre-trained model to SBEM data with staining differences and novel specific staining features (cellular nuclei) and with similar voxel size (5.9×5.9×40 nm). As expected applying the network without further refinement on the new dataset lead to unsatisfactory results (**Fig. 2d** (upper middle)). Adapting the histogram to better match the earlier dataset and applying Gaussian de-noising to increase similarity between the two image datasets was insufficient to remove the ambiguity of the prediction map introduced by new features (nucleus) and staining differences. However, we found that re-training the model with a short amount of training time (l/10^th^ of the original training time) and substantially smaller data size (l/5^th^) was sufficient to remove artifacts in the image segmentation. Thus, we used only 20 training images and labels (1024*1024 pixel) instead of 100 and were able to adapt the network with 2000 or fewer iterations to the new dataset (from 22000 to 24000 iterations for 5fm; **Fig. 2d**; 14454 to 15757 for 3fm; and 16000 to 18000 for 1fm, **Supplementary Fig. 2**). To distinguish nuclear membranes from the cellular membranes, we performed separate training for the nucleus to distinguish the nuclei from the cell somata (**Fig. 2d**). We hope that using existing models to greatly reduce the effort required for training new models will encourage the community to share their trained models and training data after publication. This could decrease the expense for image segmentation by up to 90% (**Fig. 3**), making this an appealing approach even for smaller segmentation tasks or when other image processing strategies may still be possible. For users not deeply familiar with image segmentation operations, generating training data is straight-forward (see supplementary material) and can be minimized for similar datasets.

**Figure 3:**
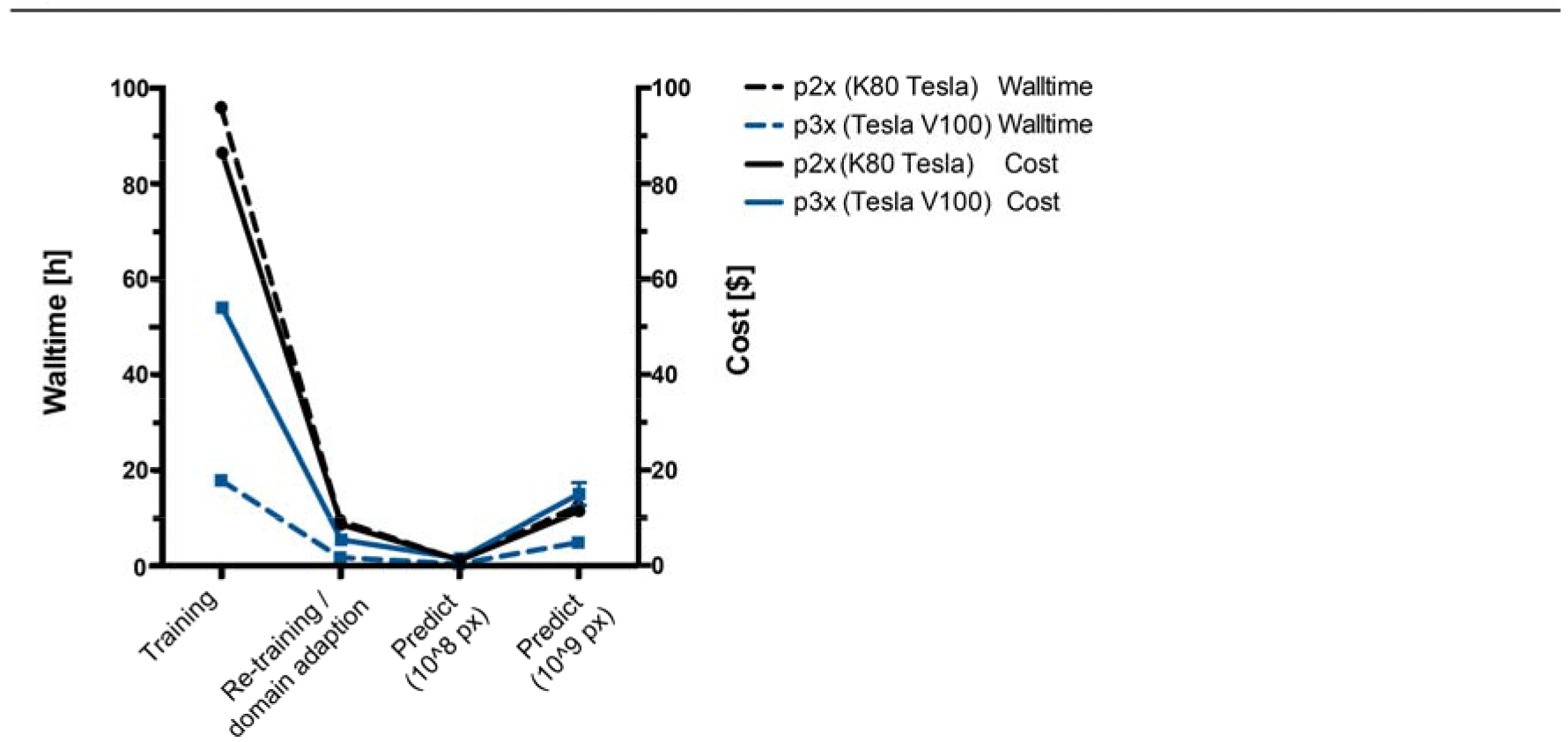
Time and cost evaluation. Training expense can be reduced substantially (to l/10^th^) by performing domain adaption from a pre-trained model (see **Fig. 2d** upper row). The time for prediction scales linearly to the size of the imaging data. Time for prediction and training both decrease with newer GPUs (Tesla V100 vs. Tesla K80). Cost calculations are based on current rates at $0.9/h for K80 and $3.06/h for V100 instances.

## Discussion

To date, the prospect of paying per hour for compute devices might at first be less appealing to members of the biomedical research community than the traditional one-time investment approach. However, the requisite high-performance computers come at a high entry price level. These resources are also outdated quickly and the maintenance of soft- and hardware will, from our own experience, make cloud computing a more cost and time effective option for most laboratories. We have further demonstrated ways to minimize costs for training and expect prices for compute time to drop rapidly as newer GPU technology is developed. Using cloud resources is scalable in times of high demand within the same laboratory and free of cost and maintenance when unused. Our cloud-based solution to provide a public AWS image is efficient for end-users, minimizing time spent for software / hardware configuration, and alleviates the burden on algorithm developers to support a community with a multitude of underlying systems and platforms. Furthermore, this will in the future give end-users immediate access to any new release. Here, we demonstrate a flexible design and ease of use that we expect will make the current release of *Deep3M* a useful tool for a wide range of applications in cell biology.

### Data and software availability

*CDeep3M* source code and documentation are available for download on GitHub (https://eithub.com/CRBS/cdeep3m) and is free for non-profit use. Amazon AWS cloudformation templates are available with each release enabling easy deployment of *CDeep3M* for AWS cloud compute infrastructure. For the end user ^~^10 minutes after creating the CloudFormation stack, a p2x or p3x instance with a fully installed version of *CDeep3M* will be available to process data. Example data are available on GitHub and trained models are available on Amazon S3. Further data will be made available upon request.

## Author Contributions

M.G.H. and M.H.E. conceived and designed the project. M.G.H., C.C. and L.T. and M.M. wrote code and analysed data. M.G.H., D.B., S. P., E.A.B., T.D. performed experiments and acquired images. M.G.H. and R.A. annotated training data. M.G.H., C.C., S.T.P. and M.H.E. wrote the manuscript with feedback from all authors.

## Acknowledgments

We thank Tao Zen for making DeepEM3D publicly available and for initial discussion. We thank Dr. Silvia Viana da Silva for critical feedback on the manuscript. We thank Sung Yeon, Nicolette M Allaway and Cesar Nava-Gonzales for help with ground truth segmentations. Research published in this manuscript has received financial support from NIH grants 5P41GM103412-29 (NCMIR), 5P41GM103426-24 (NBCR) and 5R01GM082949-10 (Cell Image Libarary (CIL). M.G.H. was supported by a postdoctoral fellowship from the interdisciplinary seed program at UCSD to build a collaborative effort to map cells, called the Visible Molecular Cell Consortium. This research benefitted from the use of credits from the National Institutes of Health (NIH) Cloud Credits Model Pilot, a component of the NIH Big Data to Knowledge (BD2K) program.

## References

1. Chen, B. C. et al. Lattice light-sheet microscopy: Imaging molecules to embryos at high spatiotemporal resolution. Science 346, 6208 (2014).

2. Bock, D. D.et al. Network anatomy and in vivo physiology of visual cortical neurons. Nature 471,177–184 (2011).

3. Long, J., Shelhamer, E. & Darrell, T. Fully convolutional networks for semantic segmentation. Proc. IEEE Comput. Soc. Conf. Comput. Vis. Pattern Recognit. 07-12-June, 3431–3440 (2015).

4. Briggman, K. L., Helmstaedter, M. & Denk, W. Wiring specificity in the direction-selectivity circuit of the retina. Nature 471,183–190 (2011).

5. Dorkenwald, S. et al. Automated synaptic connectivity inference for volume electron microscopy. Nat. Methods 14, 435–442 (2017).

6. Zeng, T., Wu, B. & Ji, S. DeepEM3D: approaching human-level performance on 3D anisotropic EM image segmentation. Bioinformatics d, 2555–2562 (2017).

7. Januszewski, M. et al. Flood-Filling Networks. 1–11 (2016).

8. Lee, K., Zung, J., Li, P., Jain, V. & Seung, H. S. Superhuman Accuracy on the SNEMI3D Connectomics Challenge. 1–11 (2017).

9. Jia, Y. et al. Caffe: Convolutional Architecture for Fast Feature Embedding. (2014). doi: 10.1145/2647868.2654889

10. Pan, S. J. & Yang, Q. A survey on transfer learning. IEEE Trans. Knowl. Data Eng. 22, 1345–1359 (2010).

11. Kasthuri, N. et al. Saturated Reconstruction of a Volume of Neocortex. Cell 162, 648–661(2015).

